# Drivers of ESBL-producing *Escherichia coli* dynamics in calf fattening farms: a modelling study

**DOI:** 10.1101/2020.09.14.296186

**Authors:** Jonathan Bastard, Marisa Haenni, Emilie Gay, Philippe Glaser, Jean-Yves Madec, Laura Temime, Lulla Opatowski

## Abstract

The contribution of bacteria in livestock to the global burden of antimicrobial resistance raises concerns worldwide. However, the dynamics of selection and diffusion of antimicrobial resistance in farm animals are not fully understood. Here, we used veal calf fattening farms as a model system, as they are a known reservoir of Extended Spectrum β-Lactamase-producing *Escherichia coli* (ESBL-EC). Longitudinal data of ESBL-EC carriage and antimicrobial use (AMU) were collected from three veal calf farms during the entire fattening process. We developed 18 agent-based mechanistic models to assess different hypotheses regarding the main drivers of ESBL-EC dynamics in calves. The models were independently fitted to the longitudinal data using Markov Chain Monte Carlo and the best model was selected. Within-farm transmission between individuals and sporadic events of contamination were found to drive ESBL-EC dynamics on farms. In the absence of AMU, the median carriage duration of ESBL-EC was estimated to be 19.6 days (95% credible interval: [12.7; 33.3]). In the best model, AMU was found to influence ESBL-EC dynamics, by affecting ESBL-EC clearance rather than acquisition. This effect of AMU was estimated to decrease gradually after the end of exposure and to disappear after 62.5 days [50.0; 76.9]. Moreover, using a simulation study, we quantified the efficacy of ESBL-EC mitigation strategies. Decreasing ESBL-EC prevalence by 50% on arrival at the fattening farm reduced prevalence at slaughter age by 33.3%. Completely eliminating the use of selective antibiotics had a strong effect on average ESBL-EC prevalence (relative reduction of 79.6%), but the effect was mild if this use was only decreased by 50% compared to baseline (relative reduction of 3.7%).

## Introduction

The detection of antibiotic-resistant bacteria in livestock animals has been a rising concern worldwide [1]. Extended Spectrum β-Lactamase (ESBL)-producing *Enterobacterales*, such as ESBL-producing *E. coli* (ESBL-EC), are a typical example as they are frequently reported in food-producing animals [2], notably calves [3–6]. These bacteria have acquired resistances to most β-lactams and are responsible for severe infections in humans [7]. The importance of considering antimicrobial resistance (AMR), and ESBL-EC in particular, in a One Health perspective is now widely recognized, due to their capability to spread across human, animal and environmental sectors [8, 9].

The drivers of AMR spread in livestock are not fully understood, although extensive antimicrobial use (AMU) is assumed to play a major role. Previous studies have investigated the relationship between variations in AMU and AMR in livestock at a scale ranging from an entire country [10–12] to specific farms [13–17], including cattle farms [18–22]. Some of these studies found an association between AMU and AMR, but not all of them. The reason may be that AMR prevalence on a farm not only depends on levels of exposure to antibiotics, but also relies upon several other factors, such as importation of animals colonised with antibiotic-resistant bacteria, within-farm transmission of antibiotic-resistant bacteria between animals and/or from humans, and contamination of animals from the environment. Moreover, antibiotic-resistant bacteria carriage and transmission are dynamic phenomena and may therefore not be well captured by classical statistical models.

Mechanistic dynamic models are useful to better understand the spread of AMR in populations [23]. They have been used extensively to study AMR spread in human populations and to assess the effect of control measures [24]. However, dynamic models simulating the transmission of AMR within farms and fitted using real longitudinal data are scarce [25–29].

Here, we propose what is, to our knowledge, the first dynamic model of AMR spread among veal calves, informed by longitudinal data on ESBL-EC carriage and AMU. Using this model, we quantitatively assess the efficacy of two different strategies to mitigate ESBL-EC prevalence on farms: decreasing ESBL-EC carriage upon arrival and decreasing AMU on fattening farms.

## Methods

### Ethics

Animal ethics approval is not required in France for rectal swabbing in calves since this is considered a non-invasive procedure.

### Study design and data collection

The field study was led between October 2015 and March 2016 in three veal calf fattening farms located in the Brittany region (France), and referred to as farms A, B and C. As a general scheme in the veal calves industry in France, fattening farms usually rear batches of 250-300 dairy calves from 3-5 weeks to 5-6 months of age before slaughter. In accordance with European animal health and welfare directives 91/629/EC and 97/2/EC, calves were kept in individual pens until eight weeks of age, and then gathered in pens housing five calves until their slaughter age. During the fattening cycle, no new calf entered the farm.

The study design is described in [30]. In brief, within each participating farm, 50 calves of the same batch were randomly tested on arrival for ESBL-EC carriage. Swabs were streaked on selective ChromID ESBL agar (bioMérieux, Marcy l’Etoile, France) and, on each farm, the 50 calves were assigned to a positive or negative ESBL-EC status based on colony growth after 24 hours at 37°C. Antimicrobial susceptibility testing was performed using the disc diffusion method and ESBL production was confirmed by the double-disc synergy test. Among these, 15 calves per farm were followed longitudinally, resulting in 45 calves in total included in the study. On each farm, calves were allocated to three different pens (five calves per pen) from eight weeks of age, and according to their initial ESBL-EC status. On farms B and C, one pen gathered calves that were all initially ESBL-EC negative, while the other two pens gathered initially ESBL-EC positive calves. On farm A, all three pens gathered ESBL-EC positive calves because all calves tested on this farm were initially ESBL-EC positive (Supplementary Material SM1).

On all three farms, rectal swabs were then collected every two weeks from each calf on days 7, 21, 35, 49, 63, 77, 91, 106, 119, 133 and 147 after the calves’ arrival, plus on day 161 for farms A and B. In total, 180 samples were collected on farms A and B, and 158 samples on farm C (SM1).

Over the study period, the antibiotics used were systematically recorded on a daily basis. Antibiotic treatments were independent from calves’ ESBL-EC carriage.

### Dynamic model

To unravel mechanisms underlying the temporal spread of ESBL-EC among calves, we built 18 variants of an agent-based discrete-time dynamic stochastic model of ESBL-EC acquisition and transmission within a calf farm. These variants included a various number of mechanisms, as described in Table 1.

**Table 1.** Description of the mechanisms included in each model variant. A full mathematical description of the models is given in the SM2.

|  | Baseline<br>constant<br>acquisition<br>rate | One<br>transmission<br>rate (within<br>farm) | Two<br>transmission<br>rates on farm<br>(within pen<br>and between<br>pens) | Effect of<br>antibiotic<br>exposure on<br>the<br>acquisition<br>rate | Effect of<br>antibiotic<br>exposure<br>on the<br>clearance<br>rate | Sporadic<br>events of<br>contamination |
| --- | --- | --- | --- | --- | --- | --- |
| Model 0 | X |  |  |  |  | X |
| Model 1 |  | X |  |  |  | X |
| Model 2 |  |  | X |  |  | X |
| Model 3 |  | X |  | X |  | X |
| Model 4 |  |  | X | X |  | X |
| Model 5 |  | X |  |  | X | X |
| Model 6 |  |  | X |  | X | X |
| Model 7 | X |  |  | X |  | X |
| Model 8 | X |  |  |  | X | X |
| Model 10 | X |  |  |  |  |  |
| Model 11 |  | X |  |  |  |  |
| Model 12 |  |  | X |  |  |  |
| Model 13 |  | X |  | X |  |  |
| Model 14 |  |  | X | X |  |  |
| Model 15 |  | X |  |  | X |  |
| Model 16 |  |  | X |  | X |  |
| Model 17 | X |  |  | X |  |  |
| Model 18 | X |  |  |  | X |  |

In the models, at each time step *t* (day), each calf was classified as either carrier or non-carrier of ESBL-EC. Model parameters are summarized in Table 2 and the full description of the models, including equations, is provided in the SM2. Some parameters were farm-specific, while the others were common to all farms (Table 2). Models were run from day 1 (arrival of calves on the farm) to day 161 (last sampling date).

**Table 2.**
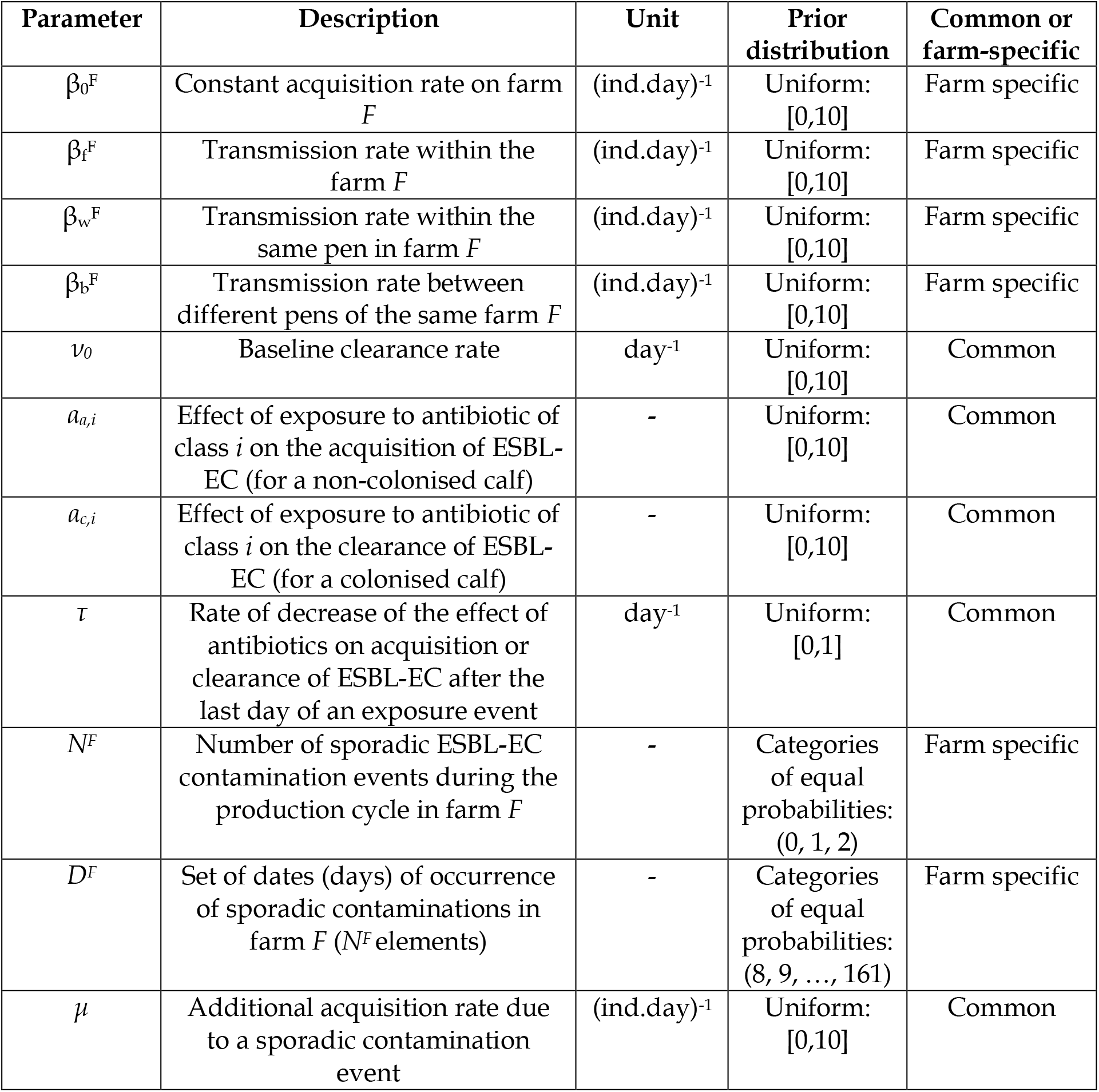
Parameters used in the dynamic models: symbol, description, unit, prior distribution and whether the parameter was farm specific (i.e. a value was estimated for each farm) or common to all farms.

#### Initialisation

The carriage status of each calf on the first day was known from the study design described above.

#### ESBL-EC acquisition

At each time *t*, the probability for an ESBL-EC negative calf to acquire ESBL-EC depended on the model variant (Table 1 and SM2). This acquisition could result either from transmission from other colonised calves, or from sporadic contaminations, depicting the possible acquisition of ESBL-EC by the calves on some specific days (estimated in the models) from another unknown source. Transmission was assumed to occur either homogeneously between calves of the same farm *F*, with rate β_f_^F^, or between calves depending on their allocated pen, assuming two transmission rates, within (β_w_^F^) and between (β_b_^F^) pens of a farm *F*. As a null hypothesis, we also investigated models which did not include any transmission between calves, but instead a constant, farm-specific, ESBL-EC acquisition rate β_0_^F^.

#### ESBL-EC clearance

At each time *t*, the probability for an ESBL-EC positive calf to clear carriage depended on a natural clearance rate, *ν_0_*, inverse of the baseline carriage duration.

#### Impact of antibiotics

Depending on the model variant, AMU was either assumed to have no effect on ESBL-EC dynamics, or to impact the probability of acquisition or clearance. For a given antibiotic class *i*, its effect on acquisition (resp. clearance) was modelled by a multiplicative factor *a_a,i_* (resp. *a_c,i_*). After the end of exposure, this effect was supposed to persist [31], but decrease exponentially [32] (i.e. tend to 1) with a rate *τ*, common to all antibiotic classes.

### Estimation and model selection

Independently for each of the 18 models, parameters were estimated in a Bayesian framework, using a Markov Chain Monte Carlo (MCMC) algorithm, implemented with the R package *rjags* [33]. Models were fitted to the data from the three farms simultaneously. Non-informative uniform priors were used for all parameters (Table 2). The 18 models were compared using the Deviance Information Criterion (DIC) [34]. Details on modelling assumptions, estimation and validation are provided in the SM3 and SM4.

To assess the quality of the best model’s fit, we simulated it by sampling parameter values in the estimated posterior distributions, and compared model predictions to observed data in each farm.

### Simulating changes in farming practices

We ran a simulation study to assess the impact of changes in farming practices on the mean prevalence of ESBL-EC carriage over the fattening cycle and on the final ESBL-EC prevalence at slaughter age, in farm A. We used the best model parameterized with posterior estimates, without sporadic events of contamination.

First, we assessed the effect of exposure to “selective” antibiotics during fattening, particularly at the beginning (from day 1), as collective “starting treatments” were a common practice to manage diseases in arriving calves. Here, we defined selective antibiotics as antibiotic classes *i* for which the estimated value of *a_a,i_* (or *a_c,i_*) was significantly different from 1 in the selected model. The baseline duration of initial exposure to selective antibiotics was defined as six days, based on data from a previous representative study led in 120 French calf fattening farms [35]. Then, we simulated reductions in the duration of this initial exposure to assess their effect on ESBL-EC prevalence.

Second, we evaluated the effect of ESBL-EC prevalence on arrival at the fattening farm (on day 1). We extracted from the literature [21] the baseline value of 68% for this prevalence on arrival in France. We assumed that changes in practices on dairy farms where calves were born could decrease this prevalence on arrival, and simulated such reductions.

In all simulations, we differentiated two scenarios. In the first scenario, the only exposure to selective antibiotics was the initial exposure described above. In the second scenario, besides the initial exposure, we simulated a 10-day “mid-cycle exposure” to selective antibiotics between days 81 and 90, to mimic the treatment of diseases during fattening.

## Results

### ESBL-EC carriage and antimicrobial use over time

ESBL-EC carriage of the 45 calves followed over the fattening cycle is detailed for each sampling date in the SM1. Time changes in the proportion of ESBL-EC positive calves in each farm is depicted in Figure 1. On all three farms, the proportion of ESBL-EC positive calves was higher on the first than on the last sampling day.

**Figure 1.**
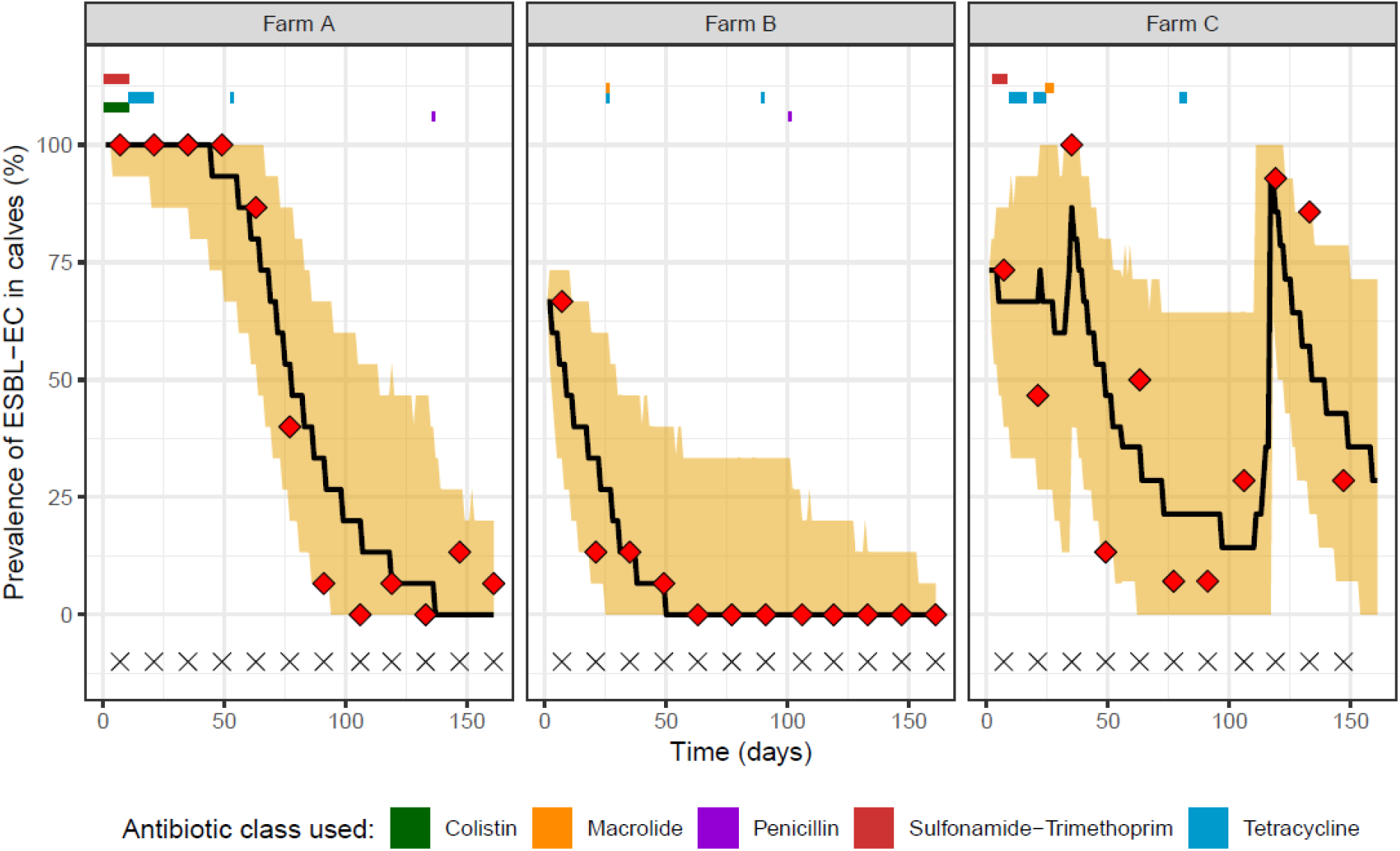
Longitudinal study of ESBL-EC colonisation in three calf fattening farms. Samples were collected every two weeks (sampling dates indicated by X) and antibiotic usage was recorded daily (period of exposure for the different classes indicated with coloured rectangles). Observed (red diamonds), median predicted ESBL-EC prevalence (black line) and 95% prediction interval for each farm, using model 5 fitted on the three farms simultaneously (1,000 repetitions of the model), are represented.

On each farm, antibiotics were always administered to all calves simultaneously over the study period, i.e. there were no individual treatments. AMU observed on farms is depicted in Figure 1 and described in the SM5.

### Parameters estimation and model comparison

Among 18 mechanistic models, model 5, which includes farm-level between-calf transmission, impact of antibiotic exposure on carriage clearance and sporadic contaminations (Table 1) presented the lowest DIC (SM6), and was therefore selected as the best model used for all analyses onwards.

The estimated posterior distributions of model 5 parameters are summarized in Table 3 and represented in the SM7. The median posterior baseline clearance rate was 0.051/day, corresponding to a median carriage duration of 19.6 days, in the absence of antibiotic exposure. The median farm-level transmission rate ranged between 0.021 and 0.059 (ind.day)^−1^, depending on the farm (Table 3).

**Table 3.**
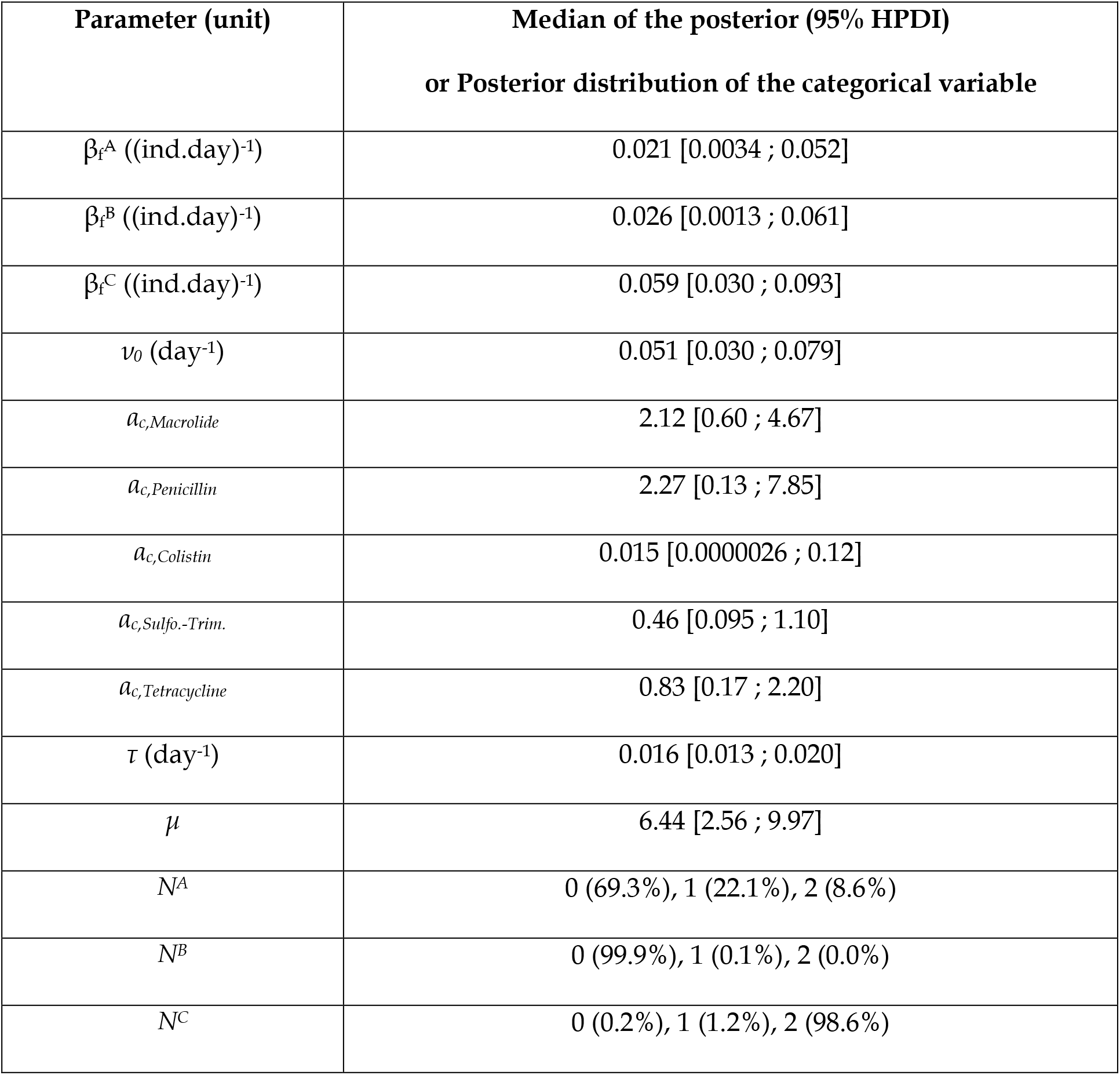
Posterior estimates of model 5 parameters: median and 95% highest posterior density interval (HPDI), or posterior distribution of the categorical variable.

Colistin exposure significantly affected ESBL-EC dynamics: being exposed to colistin on a given day was estimated to multiply the baseline clearance rate on that day by 0.015 (i.e. to divide it by 66.7) in median. Conversely, we did not find that the use of other antibiotic classes modified the baseline clearance: the 95% credibility interval included 1 for parameters *α_c,Macrolide_*, *a_c,Penicillin_*, *α_c,Sulfo.-Trim_.* and *α_c,Tetracycline_*. The effect of antibiotic exposure was estimated to decrease over time after the end of an antimicrobial use with a median rate of 0.016/day, suggesting that the antibiotics affected ESBL-EC dynamics up to 62.5 days in median after the end of exposure (Table 3 and SM8).

Most (69.3%) and almost all (99.9%) of the posterior samples in farms A and B, respectively, did not include any sporadic contamination. Conversely, there were two in farm C, at the beginning and at the end of the fattening period (Table 3 and SM7).

Model 5, estimated on the three farms combined, succeeded in fitting the observed data for each farm, as most of the observed data were in the 95% prediction interval (Figure 1). The fit of model 5 when estimated separately for each farm is shown in the SM9.

### Impact of changes in farming practices

Figure 2 represents the mean ESBL-EC prevalence over the cycle and ESBL-EC prevalence at slaughter age predicted by model 5 in farm A, when three parameters vary from their baseline values. We varied: (i) ESBL-EC prevalence in calves arriving from dairy farms, (ii) the duration of calves’ exposure to selective antibiotics on arrival (from day 1), and (iii) the duration of calves’ exposure to selective antibiotics in the middle of the production cycle (two scenarios: 0 or 10 days from mid-cycle).

**Figure 2.**
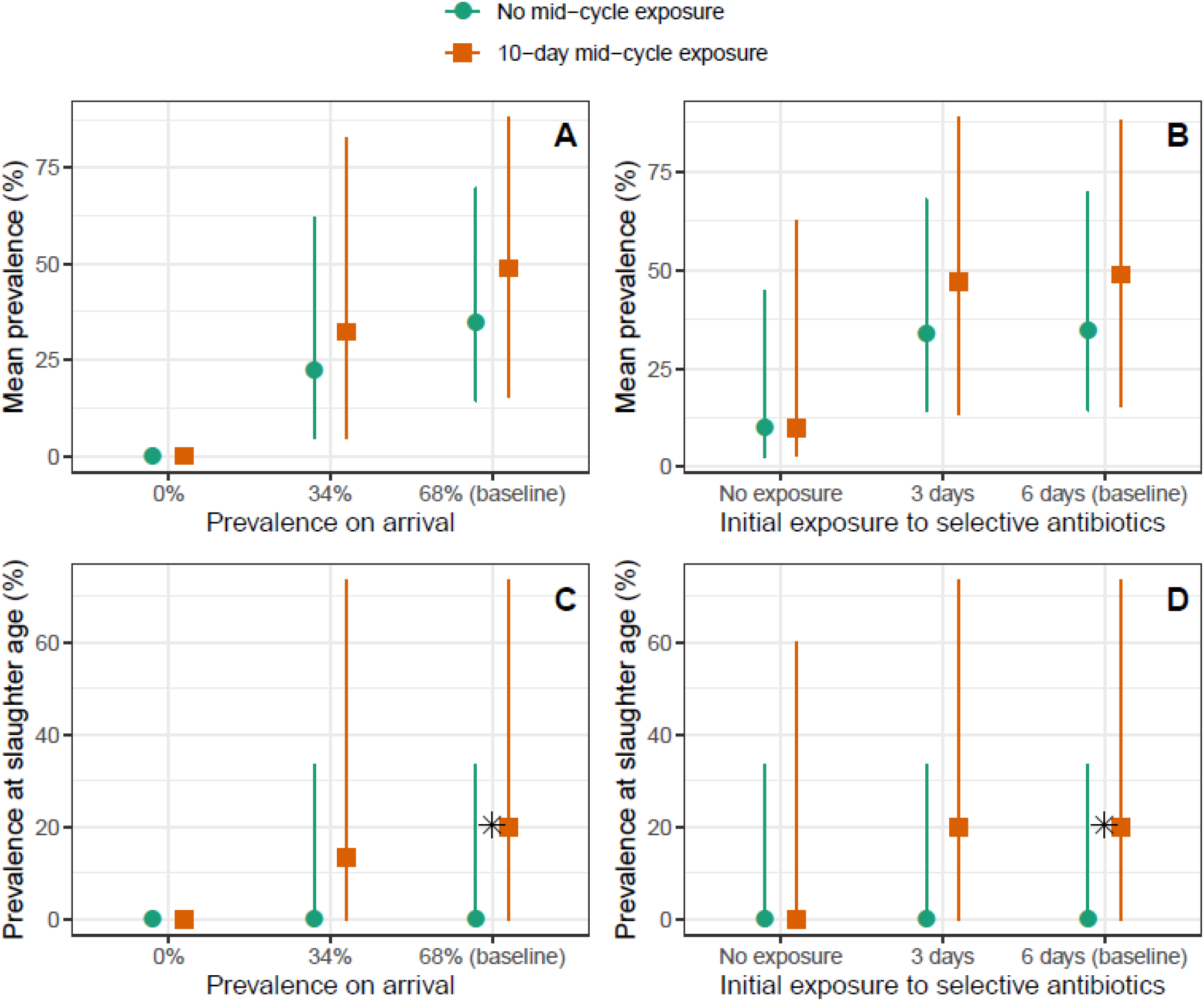
Simulations of ESBL-EC mitigation strategies. Mean ESBL-EC prevalence (panels A and B) and prevalence at slaughter age (panels C and D) predicted by model 5 (1,000 repetitions of the model), when ESBL-EC prevalence on arrival (panels A and C) and the duration of the initial antibiotic exposure (panels B and D) are changed from their baseline values. Scenarios without (turquoise) or with (orange) a 10-day antibiotic exposure in the middle of the fattening cycle (between days 81 and 90) are explored. Coloured dots are the predicted medians and intervals are 95% prediction intervals. The ✱ dot is the prevalence at slaughter age observed in [21] and is close to our baseline predictions in the scenario with a 10-day mid-cycle exposure.

In all simulations, both the mean prevalence and prevalence at slaughter age were higher in the scenario with mid-cycle exposure than without. In the baseline situation corresponding to ESBL-EC prevalence on arrival and initial exposure observed in France [21, 35], and assuming a mid-cycle selective antibiotic exposure, the predicted median ESBL-EC prevalence at slaughter age (resp. mean prevalence over the cycle) was 20.0% (resp. 48.9%), which was consistent with the 20.4% observed at slaughter age in [21] (Figure 2). Therefore, in the following, we detail results only for the scenario with mid-cycle exposure, i.e. the most conservative and realistic scenario.

If the initial exposure to selective antibiotics was completely eliminated (resp. was reduced from 6 days to 3 days), the mean prevalence over the cycle decreased by a relative 79.6% (resp. 3.7%) in median, from 48.9% to 9.96% (resp. 47.1%), and the median prevalence at slaughter age was lowered to 0 (resp. was not affected) (Figure 2B&D).

On the other hand, if ESBL-EC prevalence on arrival was cut by half compared to the baseline, from 68% to 34%, the median prevalence at slaughter age (resp. mean prevalence over the cycle) was reduced by a relative 33.3%, from 20.0% to 13.3% (resp. a relative 33.9%, from 48.9% to 32.3%) (Figure 2A&C).

## Discussion

In this study, using longitudinal data and a dynamic model, we quantitatively estimated the between-calves transmission of ESBL-EC within farms and found a significant and persistent impact of antibiotic exposure on ESBL-EC clearance. From a simulation study, we underlined the potential impact of reductions in antimicrobial use and in ESBL-EC carriage in calves arriving from dairy farms.

Consistently with previously reported dynamics in France and the Netherlands [21, 36], ESBL-EC carriage decreased from arrival to departure in all three farms (Figure 1).

Depending on the farm, the transmission rate ranged between 0.021 and 0.059 (indiv.day)^−1^, possibly reflecting differences in farm infrastructure or practices. This is in line with the ESBL-EC transmission rate of 0.06/day estimated in broilers in the Netherlands [28]. Our median estimated carriage duration of 19.6 days was also consistent with previously reported values of 12 days for multidrug-resistant *Salmonella* Typhimurium in dairy cattle [37], and 26.84 days for ESBL-EC in broilers [28].

Sporadic contaminations were necessary to reproduce carriage dynamics from farm C. Such unexplained carriage increases were observed before [36]. They may reflect a contamination from the environment, companion animals, humans, or the equipment [22].

AMU patterns, with a third of treatments within the first two weeks, half of treatments by tetracycline, and a predominance of collective treatments, were similar to previous studies in French fattening calves [21, 35].

In the best model, AMU was shown to affect ESBL-EC clearance, rather than ESBL-EC acquisition. Among five classes of antibiotics used in the farms over the study period, we only detected a significant effect of colistin on ESBL-EC dynamics, maybe due to a lack of power for other antibiotics. For instance, the wide credible interval found for penicillin (Table 3) reflects the fact that this class was hardly used in our study.

### Main limitations of our study and perspectives

First, we did not account for the diversity in genes conferring the ESBL phenotype and *E. coli* clones. However, robust results can be drawn from modelling phenotypic AMR data alone [38]. Moreover, a mechanistic transmission model fitted with genomic data would need more complexity, accounting for (i) the within-host spread of ESBL genes between *E. coli* clones *via* mobile genetic elements, and (ii) the spread of *E. coli* clones between hosts.

Second, we assumed that antibiotic classes had an identical effect on ESBL-EC dynamics in all calves of the three farms, whereas resistance patterns in ESBL-EC strains may present individual variations. Notably, the effect of colistin on ESBL-EC we found may be specific to farm A, with colistin-resistant ESBL-EC selected in this farm only. Further genomic investigations on the presence and possible different distributions of colistin resistance genes in ESBL-EC between farms may help clarify this positive effect of colistin use on ESBL spread.

### Potential implications of our results for AMR mitigation

We showed how changes in farming practices, resulting from the implementation of AMR mitigation strategies, may impact the ESBL-EC prevalence at slaughter age, reflecting the risk of its spread in the food chain, and its mean prevalence over the fattening cycle. The latter may also be of importance to human health because animals can contaminate their environment and zoonotic transmission to farmers might occur, as observed in other livestock productions [39, 40].

Regarding the use of selective antibiotics during fattening, we found a contrast between the strong effect that their complete suppression had on ESBL-EC prevalence, and the mild effect found when their use was only reduced by half (Figure 2). This non-linear effect may be explained by the persistent effect of selective antibiotics on the gut flora [41], even when they are administered for a short duration.

Reductions in ESBL-EC prevalence in calves arriving at the fattening farm simulated the hypothetical effect of actions taken at the dairy farm (e.g. reducing AMU in new-born calves or the use of waste milk from treated cows) or transportation levels (e.g. reducing calf-to-calf transmission risk). The impact of such reductions was found to be steadier, as a 50% reduction decreased by a third both the ESBL-EC prevalence at slaughter age and the average ESBL-EC prevalence over the cycle (Figure 2).

However, to lead to field application and policy, this assessment would need a more thorough benefit-cost analysis of veterinary, zootechnical and economical features, along different steps of the cattle industry. In particular, calves are administered antibiotics because they are particularly susceptible to various diseases that can affect their growth and cause mortality [42, 43].

Moreover, these figures correspond to a situation without sporadic contamination events, that can unexplainably and strongly affect ESBL-EC prevalence on farms, as discussed above. This is why biosecurity is of prime importance when considering measures to mitigate ESBL-EC carriage in calves.

## Acknowledgements

We sincerely thank Véronique Métayer for her technical contribution in samples management and antimicrobial susceptibility testing, and Aleksandra Kovacevic for her useful proofreading of the manuscript.

