## Supplementary material for "Drivers of ESBL-producing *Escherichia coli* dynamics in calf fattening farms: a modelling study"

### SM1. Longitudinal data of ESBL-EC carriage in calves

*Table S1. ESBL-EC carriage across time in calves. The last sampling day was day 161 in farms A and B, and day 147 in farm C. ESBL-EC positive (resp. negative) samples are represented as 1 (resp. 0).*

| | | | Sampling days | | | | | | | | | | | |
|---|---|---|---|---|---|---|---|---|---|---|---|---|---|---|
| **Farm** | **Pen** | **Calf** | **7** | **21** | **35** | **49** | **63** | **77** | **91** | **106** | **119** | **133** | **147** | **161** |
| **A** | **A-1** | **1** | 1 | 1 | 1 | 1 | 1 | 0 | 0 | 0 | 0 | 0 | 0 | 0 |
| **A** | **A-1** | **2** | 1 | 1 | 1 | 1 | 0 | 0 | 0 | 0 | 0 | 0 | 0 | 0 |
| **A** | **A-1** | **3** | 1 | 1 | 1 | 1 | 1 | 1 | 0 | 0 | 0 | 0 | 0 | 1 |
| **A** | **A-1** | **4** | 1 | 1 | 1 | 1 | 1 | 0 | 0 | 0 | 0 | 0 | 0 | 0 |
| **A** | **A-1** | **5** | 1 | 1 | 1 | 1 | 0 | 0 | 0 | 0 | 0 | 0 | 0 | 0 |
| **A** | **A-2** | **6** | 1 | 1 | 1 | 1 | 1 | 0 | 0 | 0 | 0 | 0 | 0 | 0 |
| **A** | **A-2** | **7** | 1 | 1 | 1 | 1 | 1 | 0 | 0 | 0 | 0 | 0 | 0 | 0 |
| **A** | **A-2** | **8** | 1 | 1 | 1 | 1 | 1 | 1 | 0 | 0 | 0 | 0 | 0 | 0 |
| **A** | **A-2** | **9** | 1 | 1 | 1 | 1 | 1 | 0 | 0 | 0 | 0 | 0 | 0 | 0 |
| **A** | **A-2** | **10** | 1 | 1 | 1 | 1 | 1 | 1 | 0 | 0 | 0 | 0 | 0 | 0 |
| **A** | **A-3** | **11** | 1 | 1 | 1 | 1 | 1 | 1 | 0 | 0 | 1 | 0 | 1 | 0 |
| **A** | **A-3** | **12** | 1 | 1 | 1 | 1 | 1 | 0 | 0 | 0 | 0 | 0 | 0 | 0 |
| **A** | **A-3** | **13** | 1 | 1 | 1 | 1 | 1 | 1 | 1 | 0 | 0 | 0 | 1 | 0 |
| **A** | **A-3** | **14** | 1 | 1 | 1 | 1 | 1 | 0 | 0 | 0 | 0 | 0 | 0 | 0 |
| **A** | **A-3** | **15** | 1 | 1 | 1 | 1 | 1 | 1 | 0 | 0 | 0 | 0 | 0 | 0 |
| **B** | **B-1** | **16** | 0 | 0 | 0 | 0 | 0 | 0 | 0 | 0 | 0 | 0 | 0 | 0 |
| **B** | **B-1** | **17** | 0 | 0 | 0 | 0 | 0 | 0 | 0 | 0 | 0 | 0 | 0 | 0 |
| **B** | **B-1** | **18** | 0 | 0 | 0 | 0 | 0 | 0 | 0 | 0 | 0 | 0 | 0 | 0 |
| **B** | **B-1** | **19** | 0 | 0 | 0 | 0 | 0 | 0 | 0 | 0 | 0 | 0 | 0 | 0 |
| **B** | **B-1** | **20** | 0 | 0 | 0 | 0 | 0 | 0 | 0 | 0 | 0 | 0 | 0 | 0 |

| | | | | | | | | | | | | | | |
|---|---|---|---|---|---|---|---|---|---|---|---|---|---|---|
| **B** | **B-2** | **21** | 1 | 0 | 0 | 1 | 0 | 0 | 0 | 0 | 0 | 0 | 0 | 0 |
| **B** | **B-2** | **22** | 1 | 0 | 0 | 0 | 0 | 0 | 0 | 0 | 0 | 0 | 0 | 0 |
| **B** | **B-2** | **23** | 1 | 1 | 1 | 0 | 0 | 0 | 0 | 0 | 0 | 0 | 0 | 0 |
| **B** | **B-2** | **24** | 1 | 0 | 0 | 0 | 0 | 0 | 0 | 0 | 0 | 0 | 0 | 0 |
| **B** | **B-2** | **25** | 1 | 1 | 0 | 0 | 0 | 0 | 0 | 0 | 0 | 0 | 0 | 0 |
| **B** | **B-3** | **26** | 1 | 0 | 0 | 0 | 0 | 0 | 0 | 0 | 0 | 0 | 0 | 0 |
| **B** | **B-3** | **27** | 1 | 0 | 1 | 0 | 0 | 0 | 0 | 0 | 0 | 0 | 0 | 0 |
| **B** | **B-3** | **28** | 1 | 0 | 0 | 0 | 0 | 0 | 0 | 0 | 0 | 0 | 0 | 0 |
| **B** | **B-3** | **29** | 1 | 0 | 0 | 0 | 0 | 0 | 0 | 0 | 0 | 0 | 0 | 0 |
| **B** | **B-3** | **30** | 1 | 0 | 0 | 0 | 0 | 0 | 0 | 0 | 0 | 0 | 0 | 0 |
| **C** | **C-1** | **31** | 0 | 1 | 1 | 0 | 0 | 0 | 0 | 0 | 1 | 1 | 1 | / |
| **C** | **C-1** | **32*** | 0 | 1 | 1 | 0 | / | / | / | / | / | / | / | / |
| **C** | **C-1** | **33** | 0 | 0 | 1 | 0 | 0 | 0 | 0 | 0 | 1 | 1 | 0 | / |
| **C** | **C-1** | **34** | 0 | 0 | 1 | 0 | 1 | 0 | 0 | 0 | 1 | 0 | 0 | / |
| **C** | **C-2** | **35** | 1 | 1 | 1 | 0 | 1 | 0 | 0 | 0 | 1 | 1 | 1 | / |
| **C** | **C-2** | **36** | 1 | 1 | 1 | 0 | 0 | 0 | 1 | 0 | 1 | 0 | 0 | / |
| **C** | **C-2** | **37** | 1 | 1 | 1 | 0 | 0 | 0 | 0 | 0 | 1 | 1 | 0 | / |
| **C** | **C-2** | **38** | 1 | 1 | 1 | 0 | 0 | 0 | 0 | 1 | 1 | 1 | 1 | / |
| **C** | **C-2** | **39** | 1 | 0 | 1 | 0 | 1 | 0 | 0 | 1 | 1 | 1 | 0 | / |
| **C** | **C-3** | **40** | 1 | 1 | 1 | 1 | 1 | 0 | 0 | 0 | 1 | 1 | 0 | / |
| **C** | **C-3** | **41** | 1 | 0 | 1 | 0 | 0 | 0 | 0 | 0 | 1 | 1 | 0 | / |
| **C** | **C-3** | **42** | 1 | 0 | 1 | 0 | 1 | 0 | 0 | 0 | 1 | 1 | 0 | / |
| **C** | **C-3** | **43** | 1 | 0 | 1 | 0 | 1 | 0 | 0 | 1 | 0 | 1 | 0 | / |
| **C** | **C-3** | **44** | 1 | 0 | 1 | 1 | 1 | 1 | 0 | 1 | 1 | 1 | 0 | / |
| **C** | **C-3** | **45** | 1 | 0 | 1 | 0 | 0 | 0 | 0 | 0 | 1 | 1 | 1 | / |

* Calf #32 died between the $4^{th}$ and the $5^{th}$ sampling days.

### SM2. Details on the models

At time $t$, each ESBL-EC negative calf could acquire an ESBL-EC with a probability

$$P = 1-e^{-A(t)}$$

where $A(t)$ was the acquisition rate, which formula depended on the model variant as detailed in Table S2. $A(t)$ was generally defined as:

$$A(t) = [A_{trans}(t) + A_{spor}(t)] \times \lambda(t)$$

with, at time $t$, $A_{trans}(t)$ the acquisition term related to between-calves transmission, $A_{spor}(t)$ the term related to sporadic contaminations and $\lambda(t)$ the multiplicative effect related to AMU.

*Transmission model*. Transmission was assumed to occur either homogeneously between calves of the same farm $F$, with rate $\beta_f^F$, or between calves depending on their attributed pen, assuming two transmission rates, within ($\beta_w^F$) and between ($\beta_b^F$) pens of a farm $F$. The transmission rate between individual pens was assumed to be the same as between collective pens. Transmission was supposed to be proportional to the proportion of calves carrying ESBL-EC. As a null hypothesis, we also investigated models, which did not include any transmission between calves, but instead a constant, farm-specific ESBL-EC acquisition rate $\beta_0^F$. Therefore, at time $t$:

$A_{trans}(t) = \beta_f^F \left(\frac{I}{S+I}\right)_{F,\ t-1}$ for homogeneous mixing (models 1, 3, 5, 11, 13 and 15)

$= \beta_w^F \left(\frac{I}{S+I}\right)_{in\ pen,\ t-1} + \beta_b^F \left(\frac{I}{S+I}\right)_{out\ pen,\ t-1}$ for pen-specific mixing (models 2, 4, 6, 12, 14 and 16)

$= \beta_0^F$ for the baseline acquisition rate (models, 0, 7, 8, 10, 17 and 18)

where $I$ was the number of ESBL-EC positive calves in the farm (within the pen, outside the pen or globally in the farm) and $S$ the number of ESBL-EC negative calves.

*Sporadic contamination*. We also assumed that sporadic contamination could occur, depicting the possible acquisition of ESBL-EC by the calves on farm $F$ on some specific days from another (unknown) source other than the colonised calves. To model this process, we assumed that sporadic contamination events in a given farm increased the acquisition rate $A(t)$ of all calves on the farm on the day of contamination by a factor $\mu$. We estimated $N^F$, the number of days with contamination events that occurred across the follow up, and $D^F$, the specific dates of contamination. This mechanism was activated in models 0 to 8.

We defined: $A_{spor}(t) = \mu$ if $t \in D^F$ (= 0 otherwise)

*Clearance.* ESBL-EC positive calves could clear carriage with a probability $1-e^{-C(t)}$, where $C(t)$ was the clearance rate at time $t$. In the baseline model, we assumed a natural clearance rate, $\nu_0$ (inverse of the baseline carriage duration).

*Impact of antibiotics.* We modelled the possible effect of AMU through two mechanisms. On the one hand, we assumed that AMU could affect the acquisition rate *A(t)*. For a given antibiotic class *i*, this was modelled by a multiplicative factor $\alpha_{a,i}$. After the end of exposure, this effect was supposed to persist [1], but decrease exponentially [2] (i.e. tend to 1) with a rate $\tau$, common to all antibiotic classes. This mechanism was active for models 3, 4, 7, 13, 14 and 17 (Table 1, main text).

Therefore, in these models, at time *t*,

$$\lambda(t) \quad = \prod_i \alpha_{a,i}^{max(0\,;\,1-\tau Ei(t))} \qquad (= 1 \text{ otherwise})$$

where $E_i(t)$ was the number of days since the last exposure of the calf to antibiotic class *i*.

On the other hand, we assumed that antibiotics could impact clearance by modulating the baseline clearance rate $v_0$ as follows (activated in models 5, 6, 8, 15, 16 and 18):

$$C(t) \quad = \nu_0 . \prod_i \alpha_{c,i}^{max(0\,;\,1-\tau Ei(t))} \qquad (= \nu_0 \text{ otherwise})$$

where $E_i(t)$ and $\tau$ were as defined above, and $\alpha_{c,i}$ was a multiplicative factor analogous to $\alpha_{a,i}$, representing the effect of exposure to antibiotic class *i* on the clearance rate *C(t).*

*Table S2. Expressions of the acquisition rate* A(t) *and clearance rate* C(t), *for the considered models.* $\lambda_a(t) = \prod_i \alpha_{a,i}^{\max(0\,;\,1-\tau Ei(t))}$ *and* $\lambda_c(t) = \prod_i \alpha_{c,i}^{\max(0\,;\,1-\tau Ei(t))}$.

| **Model** | **Acquisition rate *A(t)*** | **Clearance rate *C(t)*** |
|---|---|---|
| 0 | $A_{spor}(t) + \beta_0^F$ | $\nu_0$ |
| 1 | $A_{spor}(t) + \beta_f^F \left(\frac{I}{S+I}\right)_{F,\ t-1}$ | $\nu_0$ |
| 2 | $A_{spor}(t) + \beta_w^F \left(\frac{I}{S+I}\right)_{in\ pen,\ t-1} + \beta_b^F \left(\frac{I}{S+I}\right)_{out\ pen,\ t-1}$ | $\nu_0$ |
| 3 | $[A_{spor}(t) + \beta_f^F \left(\frac{I}{S+I}\right)_{F,\ t-1}] \times \lambda_a(t)$ | $\nu_0$ |
| 4 | $[A_{spor}(t) + \beta_w^F \left(\frac{I}{S+I}\right)_{in\ pen,\ t-1} + \beta_b^F \left(\frac{I}{S+I}\right)_{out\ pen,\ t-1}] \times \lambda_a(t)$ | $\nu_0$ |
| 5 | $A_{spor}(t) + \beta_f^F \left(\frac{I}{S+I}\right)_{F,\ t-1}$ | $\nu_0 \times \lambda_c(t)$ |
| 6 | $A_{spor}(t) + \beta_w^F \left(\frac{I}{S+I}\right)_{in\ pen,\ t-1} + \beta_b^F \left(\frac{I}{S+I}\right)_{out\ pen,\ t-1}$ | $\nu_0 \times \lambda_c(t)$ |
| 7 | $[A_{spor}(t) + \beta_0^F] \times \lambda_a(t)$ | $\nu_0$ |
| 8 | $A_{spor}(t) + \beta_0^F$ | $\nu_0 \times \lambda_c(t)$ |
| 10 | $\beta_0^F$ | $\nu_0$ |
| 11 | $\beta_f^F \left(\frac{I}{S+I}\right)_{F,\ t-1}$ | $\nu_0$ |
| 12 | $\beta_w^F \left(\frac{I}{S+I}\right)_{in\ pen,\ t-1} + \beta_b^F \left(\frac{I}{S+I}\right)_{out\ pen,\ t-1}$ | $\nu_0$ |
| 13 | $[\beta_f^F \left(\frac{I}{S+I}\right)_{F,\ t-1}] \times \lambda_a(t)$ | $\nu_0$ |
| 14 | $[\beta_w^F \left(\frac{I}{S+I}\right)_{in\ pen,\ t-1} + \beta_b^F \left(\frac{I}{S+I}\right)_{out\ pen,\ t-1}] \times \lambda_a(t)$ | $\nu_0$ |
| 15 | $\beta_f^F \left(\frac{I}{S+I}\right)_{F,\ t-1}$ | $\nu_0 \times \lambda_c(t)$ |
| 16 | $\beta_w^F \left(\frac{I}{S+I}\right)_{in\ pen,\ t-1} + \beta_b^F \left(\frac{I}{S+I}\right)_{out\ pen,\ t-1}$ | $\nu_0 \times \lambda_c(t)$ |
| 17 | $\beta_0^F \times \lambda_a(t)$ | $\nu_0$ |
| 18 | $\beta_0^F$ | $\nu_0 \times \lambda_c(t)$ |

**SM3. Details on modelling assumptions and on the estimation using MCMC**

We explored the space of possibilities for acquisition and clearance dates for each individual using data augmentation [3, 4]. Based on previous estimates of the time to clearance of ESBL-producing *Enterobacterales* [5-7], we assumed that, for a given calf, if two consecutive samples were positive (resp. negative) for ESBL-EC carriage, the calf was ESBL-EC positive (resp. negative) everyday between the two sampling dates. We checked that this hypothesis did not affect our estimations and validated our method by estimating the parameters from the analysis of simulated data (see SM4 of this Supplementary material).

Model parameters were estimated in a Bayesian framework, using a Markov Chain Monte Carlo (MCMC) algorithm, implemented with the R package *rjags* [8]. For each model variant, 3 chains were run for 6,000 iterations each. Depending on the model variant, from 2,000 to 10,000 values were discarded as burn-in, such that convergence of the Markov chains was reached. We sampled every 40$^{th}$ value because this was enough to avoid autocorrelation within the Markov chains. The effective sample size was at least 328 for posteriors of each model (953 for model 5, the model selected). A Gelman-Rubin statistic below 1.1 for all parameters was considered to indicate that between-chain variance was low enough compared to the within-chain variance [9]. The exception was model 4 where, if convergence of each chain was reached, the between-chain variance was still high, no matter the length of the burn-in. Non-informative uniform priors were used for all parameters (Table 2, main text). The 18 models were compared using the Deviance Information Criterion (DIC) [10]. The model for which the DIC was the lowest was selected as the best model (Table S3).

### SM4. Model validation

We simulated fake data to test our model's ability to retrieve true values of parameters. Our objective was also to assess the impact of our assumption to consider that, between two positive (resp. negative) samples, all days were considered positive (resp. negative). We repeated 1000 simulations of ESBL-EC carriage data using model 11 in one farm using the initial carriage of farm C and with the following set of true parameters: $\beta_f$ = 0.058 $(\text{indiv.day})^{-1}$ ; $\nu_0$ = 0.050 $\text{day}^{-1}$. As the model was agent-based and stochastic, we could not use only one "true" simulation corresponding to the true parameters. Rather, we selected the 5 repetitions minimizing the sum of the squared difference between the prevalence in the given repetition and the average prevalence over the 1000 repetitions. The objective was to select the 5 "true" carriage datasets that were the closest of the deterministic behavior of the model using true parameters. Using each of these datasets, we applied our hypothesis described above, and fitted model 11. In Figure S1, we represent the posterior distributions estimated for each of these 5 repetitions, and the "true" values of parameters. For each of the 5 cases, we re-simulated ESBL-EC carriage to check the simulated carriage episode lengths were similar to the "true" ones. In Figure S2, the distributions of true and simulated are displayed: they were similar (p-value of the Mann-Whitney-Wilcoxon test >0.05). This indicates our hypothesis on ESBL-EC carriage did not impact our estimations.

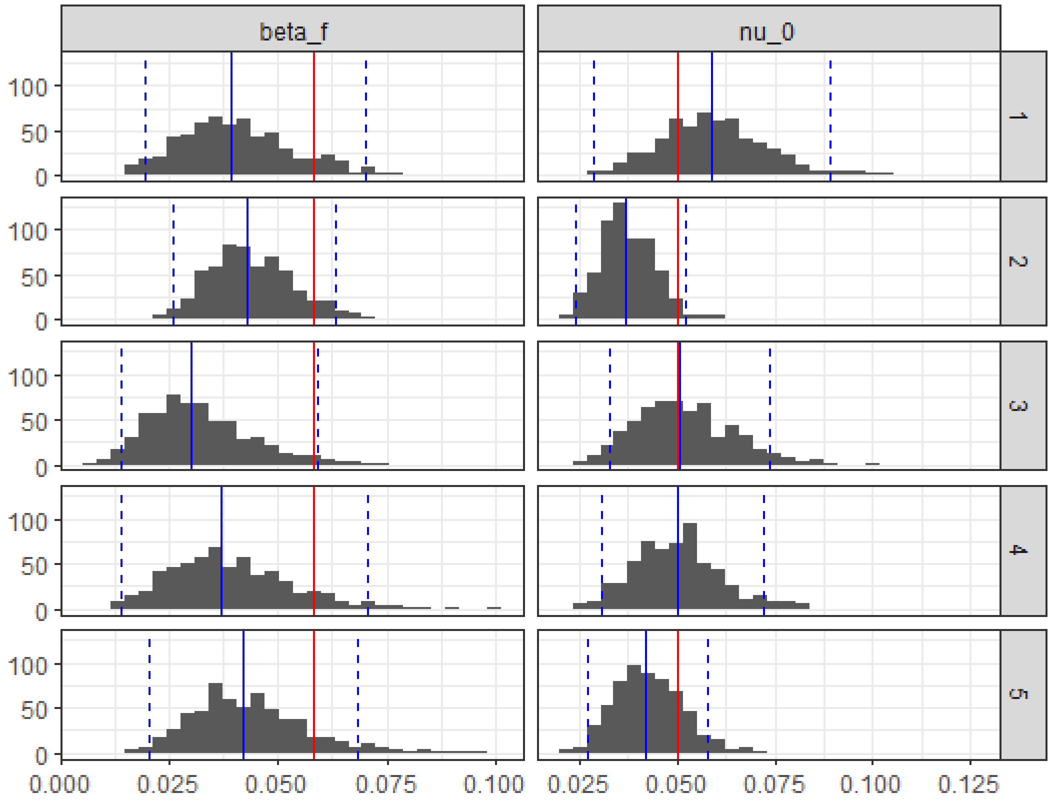


*Figure S1. For each of the 5 "true" datasets, posterior distributions (median: blue solid line ; 95% credibility interval between blue dashed lines) and true value of parameters (red solid line).*

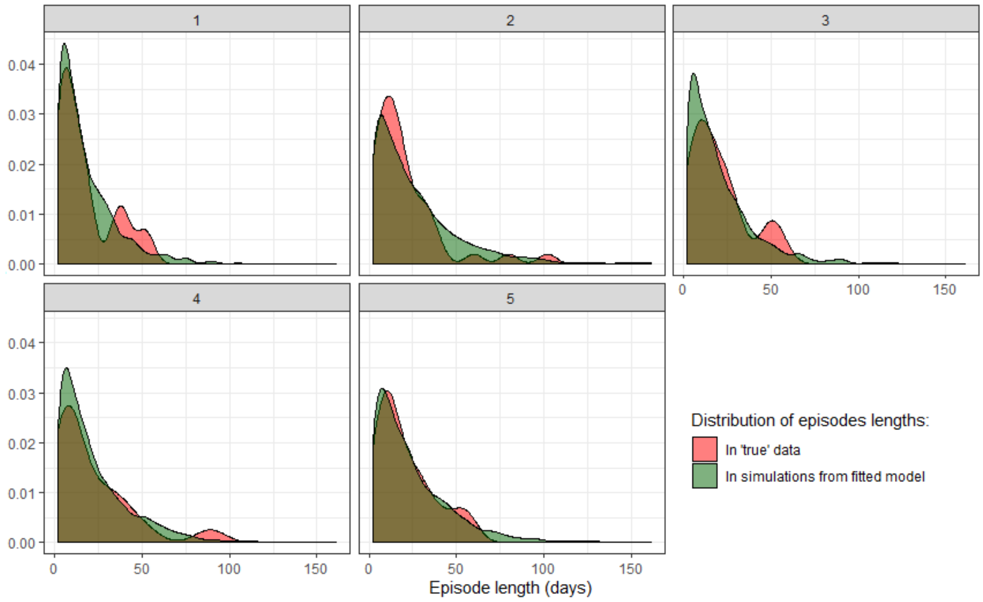


*Figure S2. For each of the 5 "true" datasets, distributions of the true ESBL-EC carriage episode lengths, and the simulated ones, after fit of Model 11.*

**SM5. Antimicrobial use observed in farms**

In our study, no individual treatment was observed: antibiotics were always administered to all calves on each farm simultaneously.

On farm A, five treatments were observed: colistin and sulfonamides between days 1 and 10, tetracycline between days 11 and 20, doxycycline on day 53, and amoxicillin on day 136 (Figure 1, main text). On farm B, calves were treated with doxycycline and erythromycin on day 26, tetracycline on day 90, and amoxicillin on day 101, resulting in four one-day treatments. On farm C, calves were exposed to five treatments: sulfonamide-trimethoprim combination between days 3 and 8, followed by tetracycline between days 10 and 16, doxycycline between days 20 and 24, spiramycin between days 25 and 27, and tetracycline between days 80 and 82.

### SM6. DIC of models

*Table S3. Deviance Information Criterion (DIC) of models. Model 5 presents the lowest DIC.*

| **Model** | **0** | **1** | **2** | **3** | **4** | **5** | **6** | **7** | **8** |
|---|---|---|---|---|---|---|---|---|---|
| **DIC** | 223.6 | 217.7 | 225.5 | 224.0 | 246.2 | 196.3 | 212.0 | 230.3 | 201.9 |
| **Model** | **10** | **11** | **12** | **13** | **14** | **15** | **16** | **17** | **18** |
| **DIC** | 256.1 | 238.5 | 248.0 | 236.6 | 250.9 | 219.1 | 232.6 | 251.6 | 239.2 |

### SM7. Posterior distributions in model 5

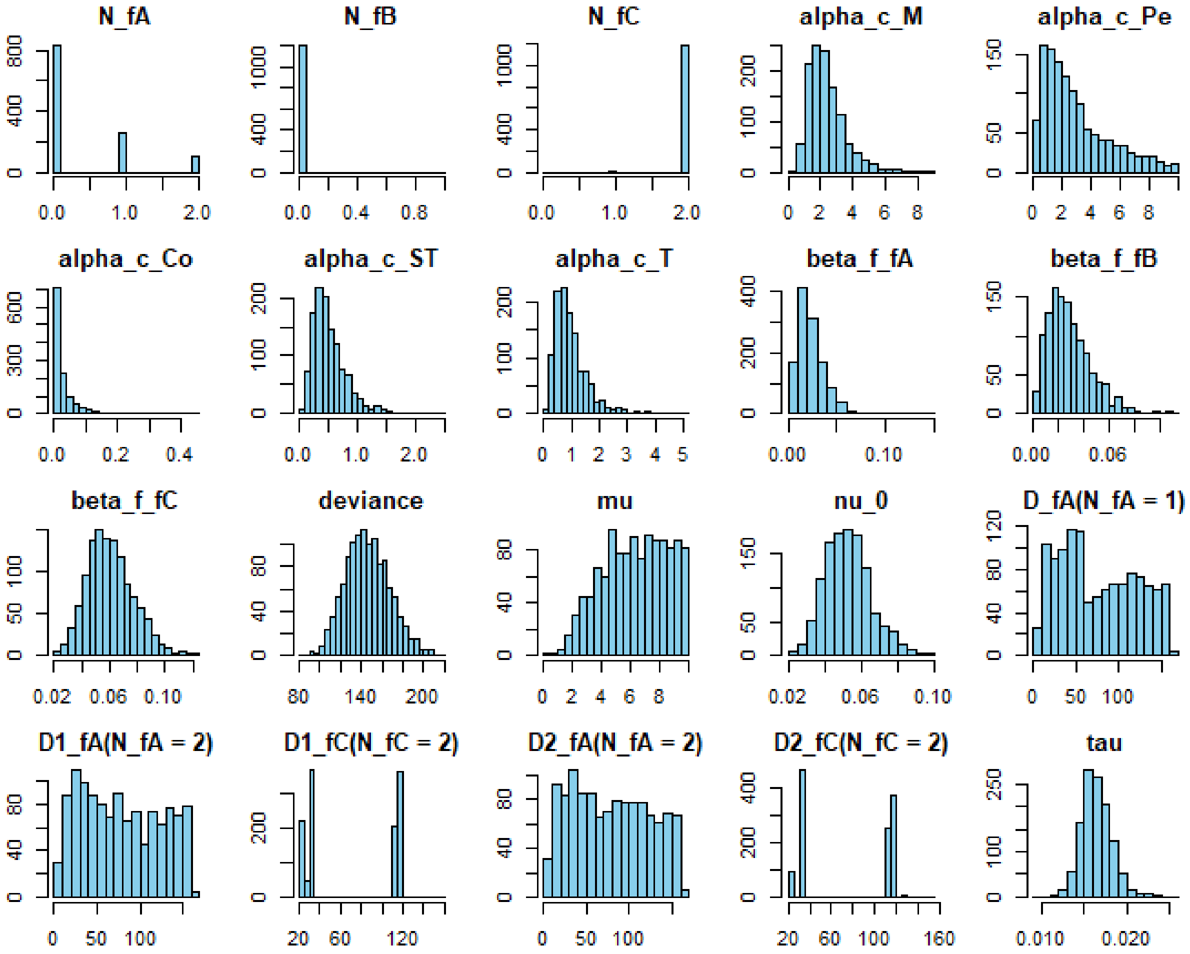


*Figure S3. Posterior distributions for the best model, model 5.*

### SM8. Evolution of the clearance rate *C(t)* after the end of colistin use

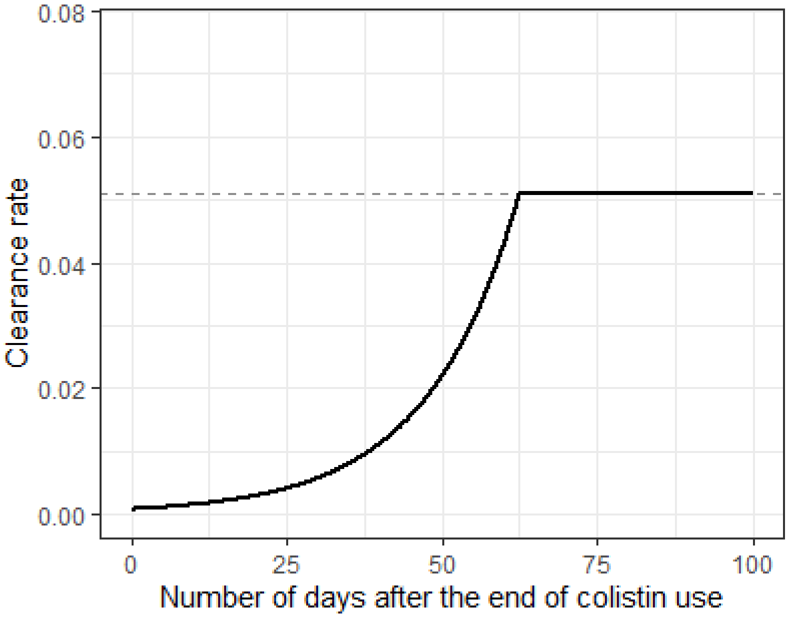


*Figure S4. Evolution of the clearance rate after the end of colistin use (solid line), using median estimated values of parameters in Model 5 (Table 3, main text). After a delay, the clearance rate* C(t) *equals* $v_0$ *(dashed line).*

**SM9. Separate fits of each farm using model 5**

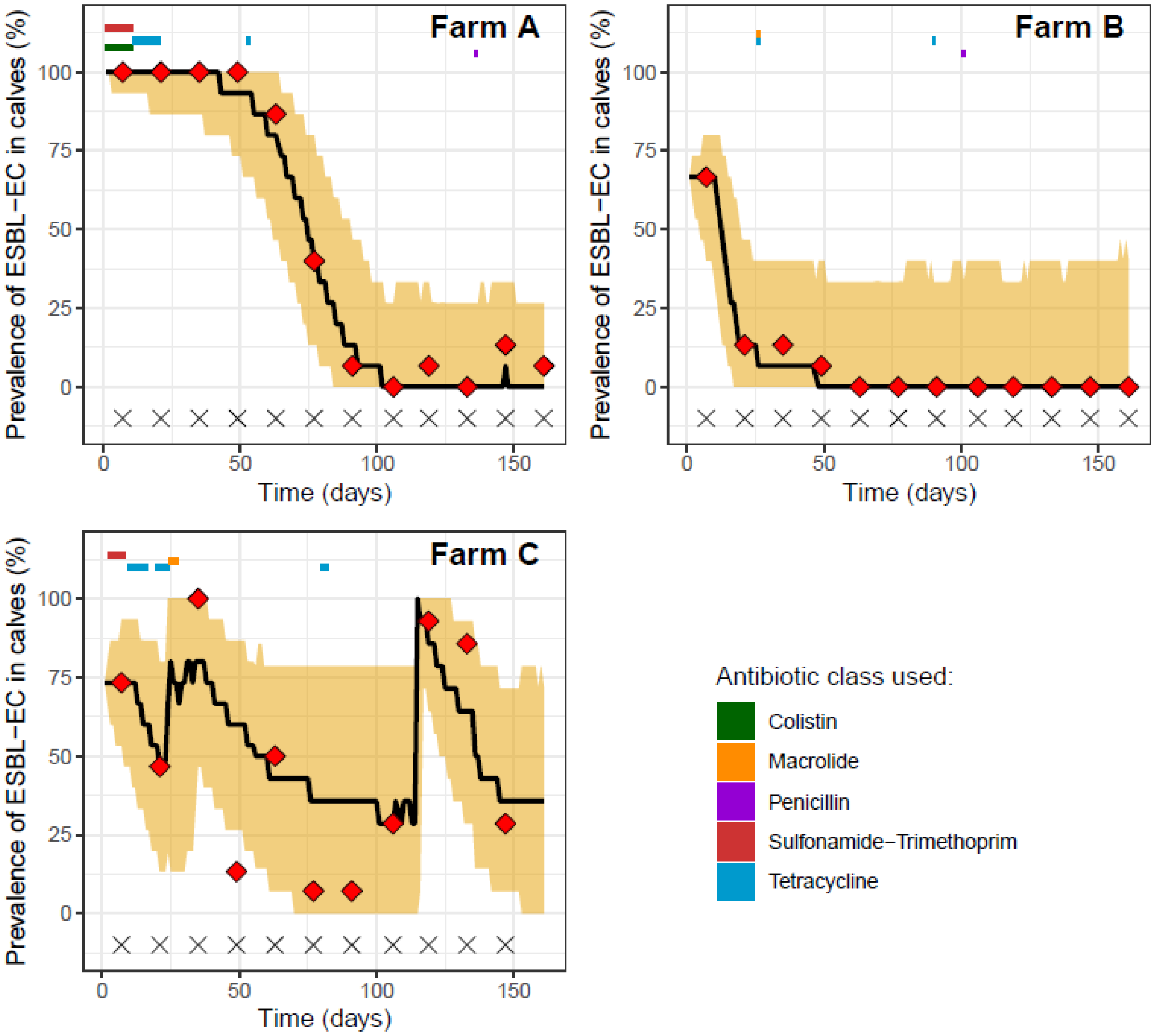


*Figure S5. Observed (red dots), median predicted ESBL-EC prevalence (black line) and 95% prediction interval using model 5 for each farm separately (1,000 repetitions of the model). Observed periods of antimicrobial use are represented for each antibiotic class.*
